# Neuroimaging: into the Multiverse

**DOI:** 10.1101/2020.10.29.359778

**Authors:** Jessica Dafflon, Pedro F. Da Costa, František Váša, Ricardo Pio Monti, Danilo Bzdok, Peter J. Hellyer, Federico Turkheimer, Jonathan Smallwood, Emily Jones, Robert Leech

## Abstract

For most neuroimaging questions the huge range of possible analytic choices leads to the possibility that conclusions from any single analytic approach may be misleading. Examples of possible choices include the motion regression approach used and smoothing and threshold factors applied during the processing pipeline. Although it is possible to perform a multiverse analysis that evaluates all possible analytic choices, this can be computationally challenging and repeated sequential analyses on the same data can compromise inferential and predictive power. Here, we establish how active learning on a low-dimensional space that captures the inter-relationships between analysis approaches can be used to efficiently approximate the whole multiverse of analyses. This approach balances the benefits of a multiverse analysis without the accompanying cost to statistical power, computational power and the integrity of inferences. We illustrate this approach with a functional MRI dataset of functional connectivity across adolescence, demonstrating how a multiverse of graph theoretic and simple pre-processing steps can be efficiently navigated using active learning. Our study shows how this approach can identify the subset of analysis techniques (i.e., pipelines) which are best able to predict participants’ ages, as well as allowing the performance of different approaches to be quantified.

## 1 Introduction

Typically, the research questions of neuroimaging studies are clearly specified (e.g., quantifying differences in functional connectivity measured with functional MRI (fMRI) between different groups or individuals), while the specific details of the analysis pipeline are not. For example, the analysis might vary in how the functional MRI data is processed to remove unwanted noise (e.g., which kernel, smoothing factor or type of motion correction to use) or what specific summary metric of functional connectivity is selected (e.g., centrality, pairwise correlation, entropy). Accordingly, it is now well recognized that a single research question can be addressed using a wide range of different analytic pipelines, often yielding slightly different answers [1, 2]. In fledgling research areas, such as functional neuroimaging, in which many of the ground truths are yet to be discovered, analytic exploration is an unavoidable aspect of the scientific process. A central conceptual question facing the community, therefore, is how to balance the data exploration needed for scientific progress with the analytical rigor necessary to minimize the number of such discoveries that are false positives. To highlight this issue [3] asked 70 independent teams to analyze the same dataset and test nine pre-defined hypothesis. Although all teams used different workflows to test these hypotheses and showed relatively high variability in the specific answers, a meta-analysis showed reasonable agreement among the broad results. The degree of consensus in studies such as this are important because different approaches can yield broadly similar answers and this in turn provides confidence that these conclusions are not tied to a specific analytic approach [4].

There is, however, a trade-off between the breadth of the initial exploratory approach and the sensitivity of subsequent scientific inferences. There are near limitless potential analysis approaches, and each additional analysis approach assessed reduces the sensitivity of statistical tests if appropriately corrected for the number of comparisons. To illustrate, if 1000 analysis approaches are tested for a specific hypothesis, then for inferential statistics it is typical to correct p-values for the multiple comparisons performed (e.g., using a Bonferroni correction); this greatly reduces the statistical sensitivity and would require the corrected p-value to be under 0.00005 to be considered statistically significant at the default p<0.05 level. Moving to out-of-sample prediction rather than inferential statistics on the whole dataset does not avoid this trade-off. The robustness of a specific conclusion from exploratory neuroimaging studies can be evaluated by training models on a specific subset of a data set and testing it on an unseen portion of the data. This approach, while minimizing problems of overfitting from yielding false positive conclusions, does not address reductions in sensitivity from performing many analyses on the same data. Here, we established a machine learning framework that maintains both exploration across analysis approaches and the sensitivity of inferential statistics and generalizability to out of sample data. Our approach allows us to explore many different features of the *universe* of pipelines and approaches, allowing many choices to be empirically compared without the need for exhaustive sampling. It does this through building a low-dimensional space across analysis approaches which is then efficiently mapped out using active learning [5].

We illustrate the utility of our multiverse approach in the context of a question in which the ground truth is completely transparent - predicting age from functional connectivity obtained from functional MRI images of adolescent and young adult participants [6]. In particular, we focus on graph theoretic analyses. The motivation for the choice of graph theoretical analysis is two-fold: (i) graph theory applied to fMRI data has been shown to be a useful technique for exploring the interrelationships between brain regions [7, 8]; (ii) such approaches have also been shown to be highly sensitive to pre-processing steps such as thresholding [9, 10]. Moreover, there are dependencies between different types of graph theory measures [11] such that the optimal analysis approach for any given dataset or question is typically unknown a priori. Indeed, previous work has demonstrated that variations in (structural) brain network construction and analysis pipelines have a substantial impact on results [12]. Focusing on the graph theoretical analysis of functional brain networks with the aim of predicting age, allows us to evaluate the utility of the multiverse approach under conditions when the ground truth is known, but the ideal analytic approach is not. We note, however, that the approach could be applied more generally to many different types of neuroimaging problems (both functional and structural) or indeed other types of data (e.g., univariate and multivariate analyses), and ultimately be applied to basic scientific and clinical research questions when the ground truth are less transparent.

## 2 Methods

### 2.1 Rationale and analytical workflow

While the proposed framework can be applied to any neuroimaging studies, here we illustrate its capabilities in the context of predicting age. The framework consists of two main steps: (i) create a low-dimensional continuous space of the different analysis approaches; (ii) an active learning component that efficiently searches the created space to find the optimal analysis pipeline (in this case, the pipeline which best predicts age) and produces estimates of other pipeline’s performance. We focused on predicting brain age as it has been proposed as a useful biomarker of neurological and psychiatric health [13, 14] and is predictive of a range of other factors, including mortality [15]. More generally, predicting age is a useful proof of principle for methodological demonstrations since the participant’s age is known with certainty [16] and a range of studies have shown that functional connectivity from resting state correlates with age (e.g., [6, 17, 18]). All code used for analyses and figure generation is available on GitHub (https://github.com/Mind-the-Pineapple/into-the-multiverse) and can be run using Colab. In the next section, we first describe the data, the range of analysis approaches considered, how the analysis space was constructed and finally, the active learning approach used to sample the space and so be able to estimate brain age prediction across the multiverse of analysis approaches without exhaustive testing.

### 2.2 Functional connectivity data

The starting point is a functional MRI dataset of changes in functional connectivity across adolescence from [6]. This dataset consists of 520 scans from 298 neurologically healthy individuals (age 14-26, mean age = 19.24, see [6] for details). Here, we only performed cross-sectional analyses and so only kept the first scan for each individual. The dataset was split in two parts: (i) 50 individuals, selected at random, were used to build the low-dimensional space; (ii) the remaining 248 individuals were subsequently used to perform search and validation on the space.

#### Analysis approaches

There are many decisions necessary to conduct a functional connectivity study, including choices regarding data acquisition, pre-processing, summary metrics and statistical models. Here, for convenience, we use already acquired data which has been through extensive pre-processing pipelines to reduce many potentially confounding sources of non-neural artefacts. Usefully, two pre-processed datasets were shared by [6] with two different types of correction for movement artefacts: (i) global signal regression and (ii) motion regression. The pre-processed fMRI time-series had been averaged within 346 regions of interest, including 330 cortical regions from the Human Connectome Project multi-modal parcellation [19] (excluding 30 "dropout" regions with low signal intensity) and 16 subcortical regions from Freesurfer. The Pearson correlation coefficient was used to calculate functional connectivity (FC) between these regions. For further details regarding data pre-processing, see [6]. In future work, the approach outlined below could be applied to all of the image pre-processing steps, the choice of parcellation and the connectivity metric for a more extensive multiverse analysis.

In order to demonstrate the capabilities of the proposed method we consider varying three distinct analysis pipeline choices. These are:

1. *The nature of regression:* we explored data from two types of data regression motion regression and global signal regression.
2. *The graph theoretical metric studied:* we consider 16 distinct metrics covering both simple and higher-level metrics which have been successfully employed in previous neuroimaging studies. Metrics were taken from the Python implementation of the Brain Connectivity Toolbox [20]; metrics included were those that produced nodal metrics. If prior community assignment information was required, we used the well-known Yeo network parcellation of the brain into 7 networks [21].
3. *The choice of threshold for estimated functional connectivity matrices:* we consider 17 distinct threshold values ranging from 0.4 (resulting in highly sparse networks) to 0.01 (resulting in dense networks).

The full space of analysis options is presented in Table 1. Every single evaluated pipeline was built by using one of the regression choices, one threshold and one graph theory metric. Therefore, the analyzed multiverse space consisted of 2 *×* 16 *×* 17 = 544 different analysis approaches.

**Table 1:** Different analysis approaches used to create the pipelines to be evaluated. Every pipeline was composed of one type of regression data, one graph theory metric and a threshold leading to the creation of 2 × 16 × 17 = 544 different analysis pipelines.

| Data | Graph Theory Metric | Threshold |
| --- | --- | --- |
| Motion regression | Degree | 0.4 |
| Global signal regression | Strength | 0.3 |
|  | Betweenness-centrality | 0.25 |
|  | Binary clustering coefficient | 0.2 |
|  | Weighted clustering coefficient | 0.175 |
|  | Eigenvector-centrality | 0.15 |
|  | Subgraph-centrality | 0.125 |
|  | Local efficiency | 0.1 |
|  | Modularity (Louvain) | 0.09 |
|  | Modularity (ProbTune) | 0.08 |
|  | Participation coefficient | 0.07 |
|  | Module Degree ZScore | 0.06 |
|  | Pagerank-centrality | 0.05 |
|  | Diversity coefficient | 0.04 |
|  | Gateway degree | 0.03 |
|  | K-core centrality | 0.02 |
|  |  | 0.01 |

### 2.3 Constructing a low-dimensional space of analysis approaches

In order to efficiently sample across a large number of analysis approaches, we need information about their general relationships. This is achieved by building a low-dimensional space that quantifies the similarity between approaches in terms of a distance in the low-dimensional space (i.e., how similar is the obtained functional connectivity between the different approaches). To ensure utility to a range of questions, the space should capture the similarity between approaches across a range of different problems (and potentially a range of different datasets, although that is not assessed here); for example, the same space can be used to ask questions about age (in the present case), but could equally be used to ask about other sources of individual variability (e.g., neuropsychiatric symptoms or cognitive ability). Below, we illustrate one approach to building such a space (precisely how this is done will vary depending on the type of data and other factors).

To construct the low-dimensional space, we applied all 544 analysis approaches to 50 randomly selected participants’ individual FC data (Fig. 1, A-C). The aim was to build a space to locate approaches in terms of how well they capture individual variability; therefore, for each approach, we calculated the Euclidean distance matrix (in terms of regional graph theory metrics) between different participants, e.g., the Euclidean distance of betweenness centrality across all 346 regions for each pair of participants (Fig. 1, D-F). These were subsequently reshaped into a 2D matrix corresponding to between-participant distances (this led to a matrix of 1225 participant pairs by 544 analysis approaches). Finally, the low-dimensional space was constructed with established embedding algorithms; we explored five different algorithms: local linear embedding [22], spectral embedding [23], t-distributed stochastic neighbor embedding (t-SNE) [24], multi-dimensional scaling (MDS) [25] and Uniform Manifold Approximation and Projection (UMAP) [26]. The objective of the embeddings was to create a space useful for active learning which would both: (i) capture similarity between approaches in terms of continuous distance in the space; as well as, (ii) distribute approaches relatively evenly across the space. Based on observations of the spaces resulting from the aforementioned embedding algorithms, MDS was selected to use in subsequent active learning (see Results).

**Figure 1:**
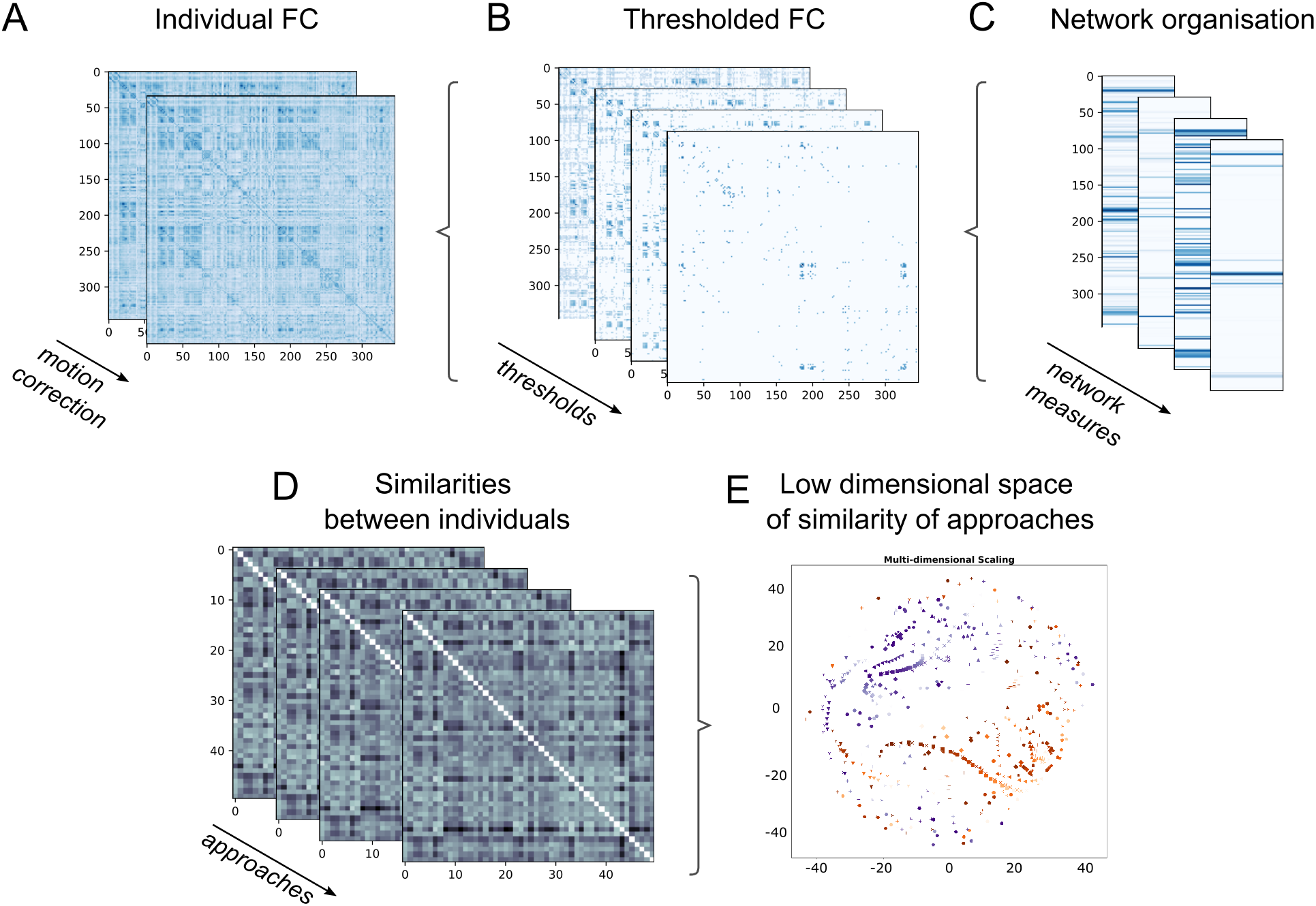
Creating a low-dimensional space to characterize the *multiverse* of different analysis approaches. A low-dimensional space was constructed using 50 (randomly selected) participants’ functional connectivity (FC) matrices. Each participants’ data was analyzed using 544 possible analysis approaches, which were composed by choosing from: two methods for motion correction (A), 17 different sparsity thresholds (B), and 16 different graph theory metrics (C). We evaluated the different approaches as how well they capture individual variability by evaluating the pairwise cosine similarity for all the different analysis approach (D). The distance matrices were then converted into a low-dimensional (2D) space summarizing the similarity between approaches by using an embedding algorithm (such as multi-dimensional scaling (E)).

### 2.4 Searching the space

Using the low-dimensional space created with FC data from 50 participants, active learning was subsequently used with the remaining 248 participants, to sparsely sample the space in order to: i) find the most successful approaches for predicting participant age based on FC; and, (ii) estimate age prediction ability for all models, including the large majority of models which were not sampled. Active sampling is performed using closed-loop Bayesian optimization with Gaussian processes [27]. This loop involves: selecting a point in the space to sample; evaluating it in terms of 5-fold cross-validated predictive accuracy; fitting a Gaussian process (GP) regression to the space; and, evaluating an acquisition function using the GP regression to select the next point to sample.

When a point in the space is selected, the closest analysis approach to that point in the space is selected and its predictive accuracy evaluated by using support vector regression for the brain age prediction. Gaussian process regression was implemented by the scikit-learn library [28] [29]) using the RBF kernel and the default parameters. It is important to highlight that although the algorithm to predict age was kept constant, the input data varied depending on the selected analysis pipeline which could have used different motion correction, thresholding or graph theory metric. Predictive accuracy was calculated with 5-fold cross-validated negative mean absolute error.

For the examples presented in the results, there was an initial *burn-in* phase in which ten points in the space were randomly selected and evaluated before active learning began. Bayesian Optimization using the upper confidence bound acquisition function [27]. The Gaussian process regression model used a Mattern kernel combined with a white noise kernel, with kernel hyperparameters chosen in each iteration by maximizing log-marginal-likelihood using the default optimizer.

## 3 Results

The first step was to construct a low-dimensional space of the analytic space. Six approaches were considered and are presented in Figure 2; in addition, results obtained with a Principal Component Analysis (PCA) are shown in Figure S1. All embedding algorithms demonstrate considerable structure in the position of the different approaches (e.g., similar types of motion correction, thresholding, graph metric are generally proximal). This suggests that the low-dimensional space captures the intended similarity between the approaches. We used a dissimilarity score to assess how much the different embedding algorithms preserved the topological information (i.e., similar analysis approaches should stay close to each other after the embedding; Figure S1). MDS, t-SNE and UMAP efficiently maintained the neighborhood of the original space. In addition, MDS displayed a relatively even spread of approaches across the whole space, especially when contrasted with Local-Linear-Embedding and Spectral Entropy. An approximately even spread across the space is desirable for the subsequent active learning and Gaussian process regression. As such, MDS was used in subsequent analyses.

**Figure 2:**
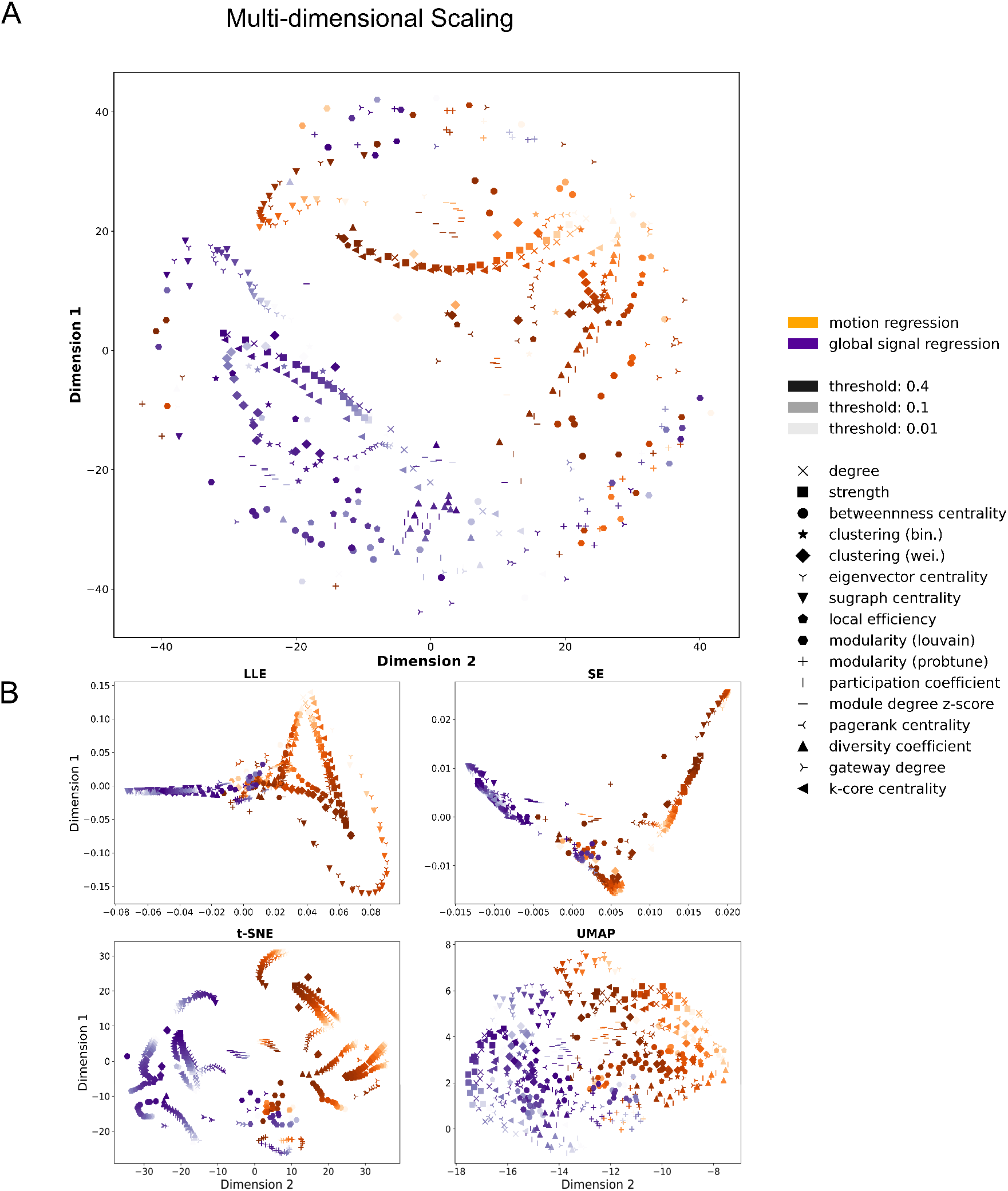
Low-dimensional embeddings of the different analysis approaches. Each point represents a combination of data accounting for noise confounds, thresholding of connectivity weights and different graph theory metrics. In particular, the colors represent both motion correction methods used to pre-process the data (i.e., motion regression (orange) and global signal regression (blue)), the color intensity represents the different thresholds used in each analysis and every graph theory metric is represented by a different symbol. A) Multi-dimensional scaling. B) Four other types of embedding: Local linear embedding (LLE), Spectral embedding (SE), t-Stochastic Neighbor Embedding (t-SNE) and Uniform Manifold Approximation and Projection (UMAP).

There are two objectives for the use of active learning on the MDS-defined space of different analysis approaches: i) finding an approximately optimal analysis approach efficiently, controlling the number of multiple comparisons; while, ii) approximately estimating performance on the multiverse of approaches without exhaustive sampling. These two objectives can be observed in Figures 3-5 where age-prediction models were trained and evaluated for different analysis approaches selected by active learning.

**Figure 3:**
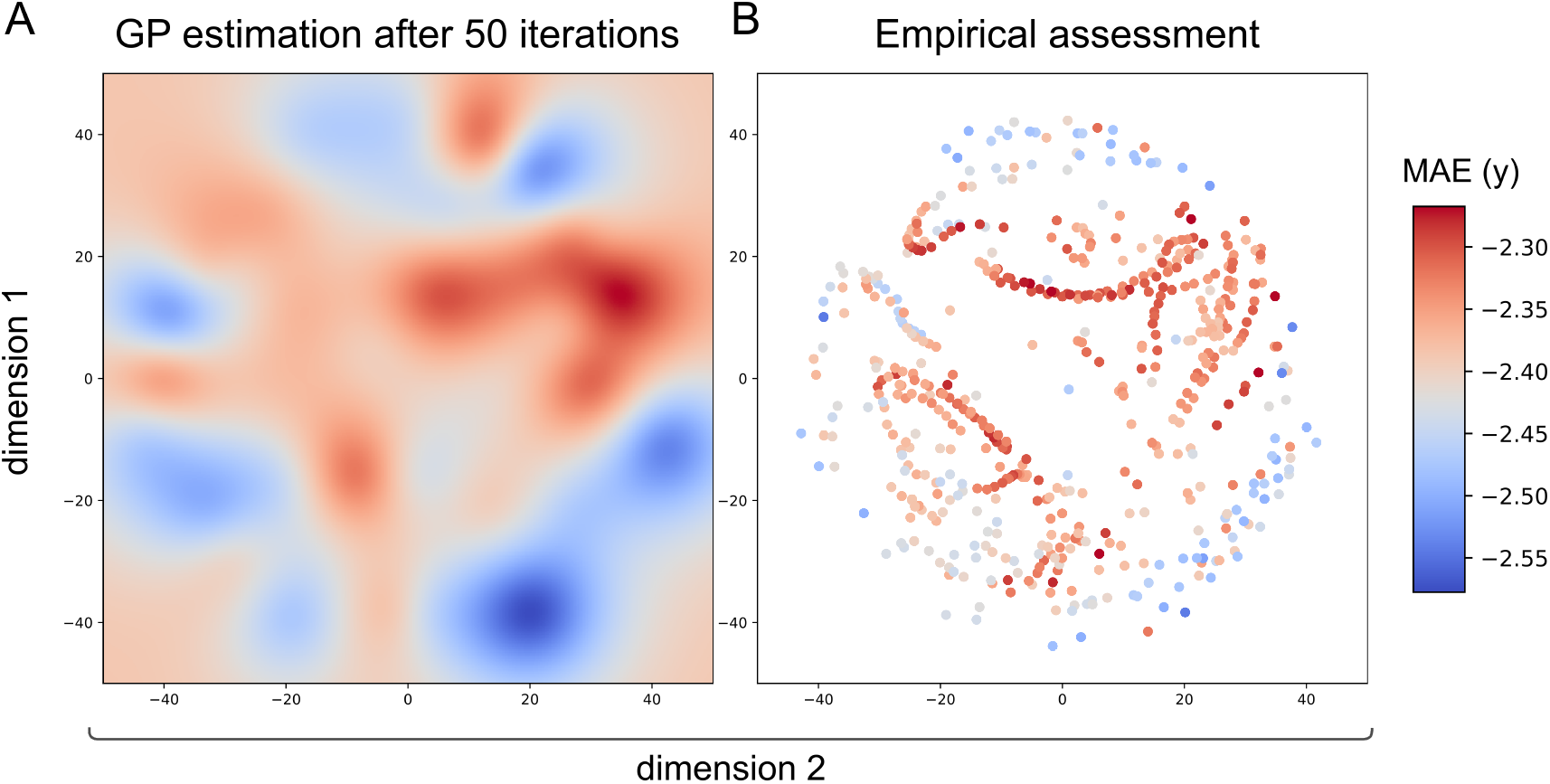
Learning the space and identification of the optimal analysis pipeline for age prediction. A) After 50 iteration of Bayesian Optimization the Gaussian Process (GP) closely estimates the empirical space. B) Empirical assessment of age prediction across the whole space. The colors correspond to negative mean absolute error of each model in years. Values closer to zero represent a more accurate prediction and are shown in red.

First, in Figure 3, we see the result of the Gaussian process regression after 50 iterations of active learning. Based on the 50 different analysis approaches sampled, GP regression estimates performance across all 544 approaches (Fig. 3A); this identifies areas predicted to have higher age-prediction performance (in warm colors) including the optimum, as well as approaches which perform worse (in cooler colors). For comparison, the ground truth of performance across the space (from exhaustive sampling of every approach) is presented in Fig. 3B. We observe a generally good concordance between actual age prediction for each approach and the estimated prediction across the whole space (Spearman’s *ρ* = 0.61, *p* < 0.0001).

The evolution of the active sampling and Gaussian process regression model is presented in Figure 4. We initially observe a poor GP estimation of the space based on the first 10 random burn-in samples. As the sampling increases, the space is progressively better estimated achieving increasingly higher correlations between empirical and estimated spaces. Acquisition function parameters strongly affect the active sampling; to illustrate this, the parameter *κ* was varied to conduct both exploratory (*κ* = 10, Fig. 4A) and exploitative versions of active sampling (*κ* = 0.1, Fig. 4B). The exploratory version achieves a better estimation of the whole space, while the exploitative version focuses on an estimated optimum much more quickly, but the GP model changes much less subsequently, resulting in a much lower correlation between estimated and empirical accuracies across the space.

**Figure 4:**
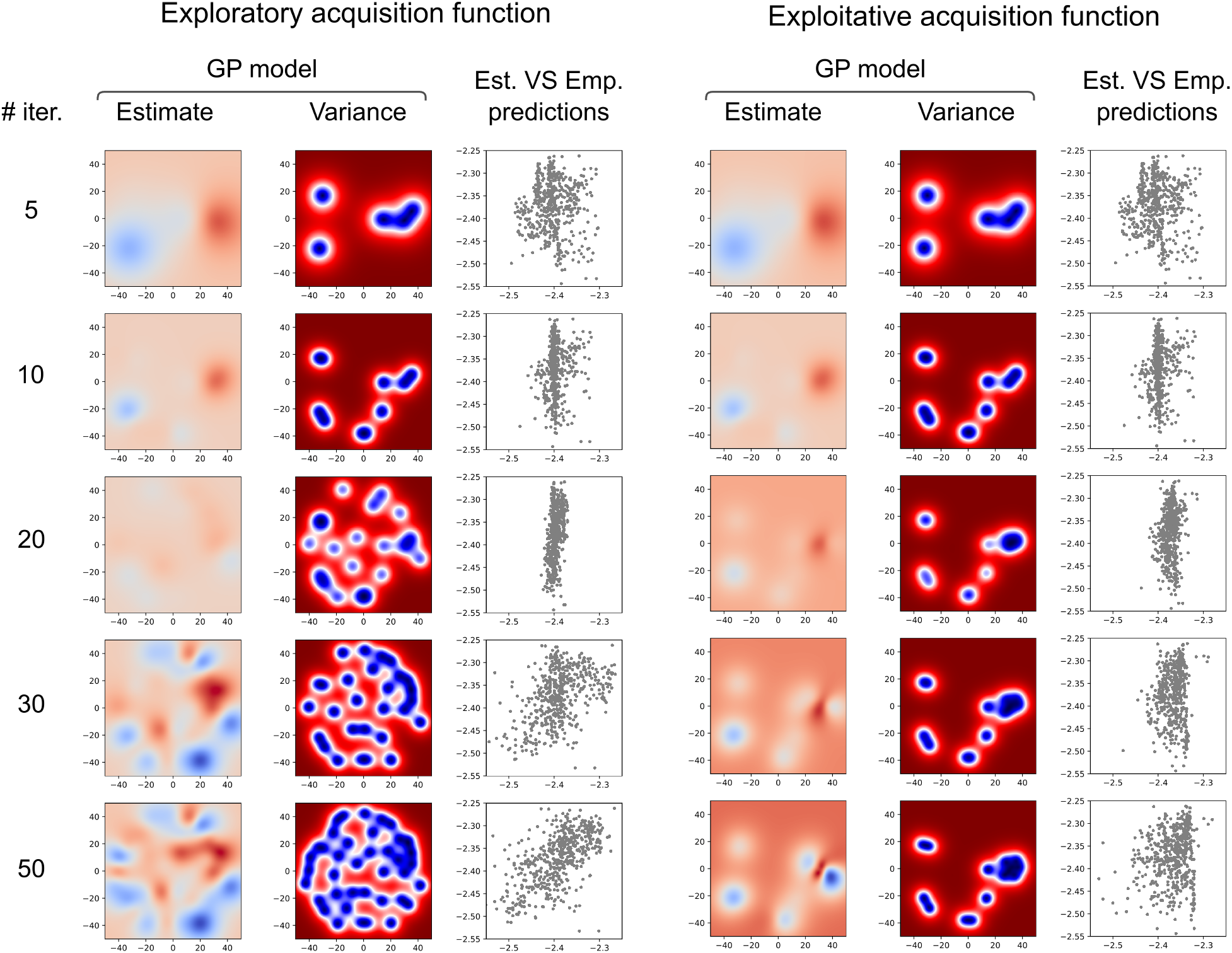
The evolution of the search across the space for: A) a more exploratory acquisition function; and, B) a more exploitative acquisition function. Within each panel, the first column is the estimated Gaussian Process (GP) model after different numbers of samples; the second column is the variance of the GP model across the space, indicating which points have been sampled; the third column is the estimated versus empirical predictions for all the analysis approaches in the space.

To investigate the reliability of the active sampling, the process was repeated 20 times (using the more exploratory *κ* = 10) with different random seeds (and so different initial random burn-in sampling). In Figure 5, the optima (i.e., model with the the highest empirical accuracy) of the 20 repetitions are represented by the black dots, based both on the highest accuracy estimated using the GP model (Fig. 5A) and for the actual sampled points (Fig. 5B). Table 2 presents the optimal analysis approaches selected by each iteration. We note that many of the optima illustrated in Table 2 were obtained by using the Betweenness centrality. This might suggest that this graph theory metric is more robust to the usage of different pre-processing choices. The range of the mean absolute error for the different optima selected versus the full range of mean absolute errors across the whole space is presented in Fig. 5C and the range of correlations between actual and estimated accuracies across the whole space for the 20 replications is presented in Fig. 5D. For inferential statistics, the optimal analysis approach selected for each of the 20 replications was assessed with a permutation test on cross-validated predictions (with 5000 random permutations), resulting in a range *p* = 0.004 − 0.044 (Bonferroni corrected for 20 samples). Note that, exhaustive sampling would result in having to correct the best model for 544 comparisons rather than 20 (requiring an uncorrected *p* < 0.000091 rather than *p* < 0.0025).

**Table 2:**
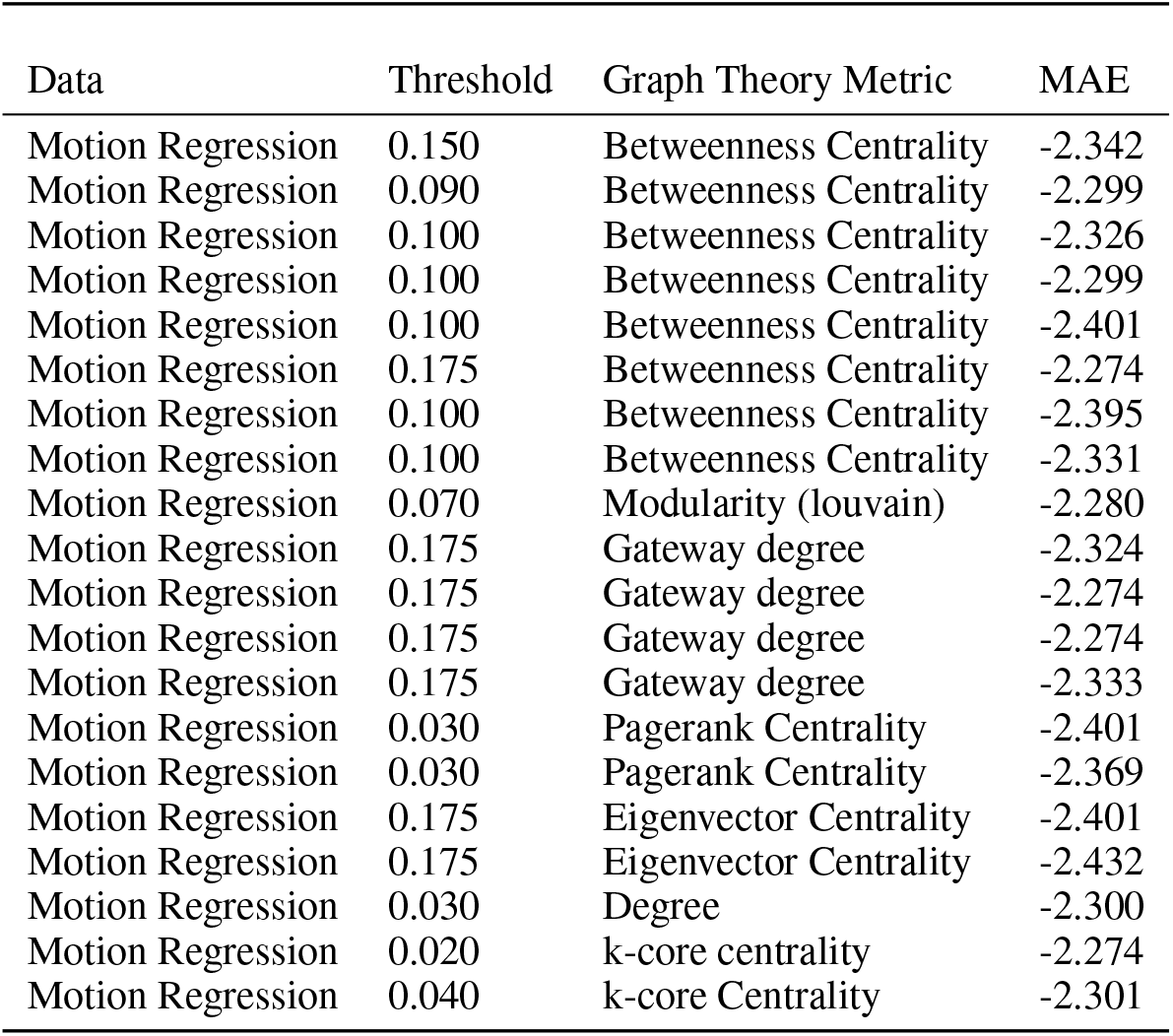
List of the data, threshold, graph theory metric and obtained mean absolute error (MAE) for the empirical optima obtained for the 20 iterations

| Data | Threshold | Graph Theory Metric | MAE |
| --- | --- | --- | --- |
| Motion Regression | 0.150 | Betweenness Centrality | -2.342 |
| Motion Regression | 0.090 | Betweenness Centrality | -2.299 |
| Motion Regression | 0.100 | Betweenness Centrality | -2.326 |
| Motion Regression | 0.100 | Betweenness Centrality | -2.299 |
| Motion Regression | 0.100 | Betweenness Centrality | -2.401 |
| Motion Regression | 0.175 | Betweenness Centrality | -2.274 |
| Motion Regression | 0.100 | Betweenness Centrality | -2.395 |
| Motion Regression | 0.100 | Betweenness Centrality | -2.331 |
| Motion Regression | 0.070 | Modularity (louvain) | -2.280 |
| Motion Regression | 0.175 | Gateway degree | -2.324 |
| Motion Regression | 0.175 | Gateway degree | -2.274 |
| Motion Regression | 0.175 | Gateway degree | -2.274 |
| Motion Regression | 0.175 | Gateway degree | -2.333 |
| Motion Regression | 0.030 | Pagerank Centrality | -2.401 |
| Motion Regression | 0.030 | Pagerank Centrality | -2.369 |
| Motion Regression | 0.175 | Eigenvector Centrality | -2.401 |
| Motion Regression | 0.175 | Eigenvector Centrality | -2.432 |
| Motion Regression | 0.030 | Degree | -2.300 |
| Motion Regression | 0.020 | k-core centrality | -2.274 |
| Motion Regression | 0.040 | k-core Centrality | -2.301 |

**Figure 5:**
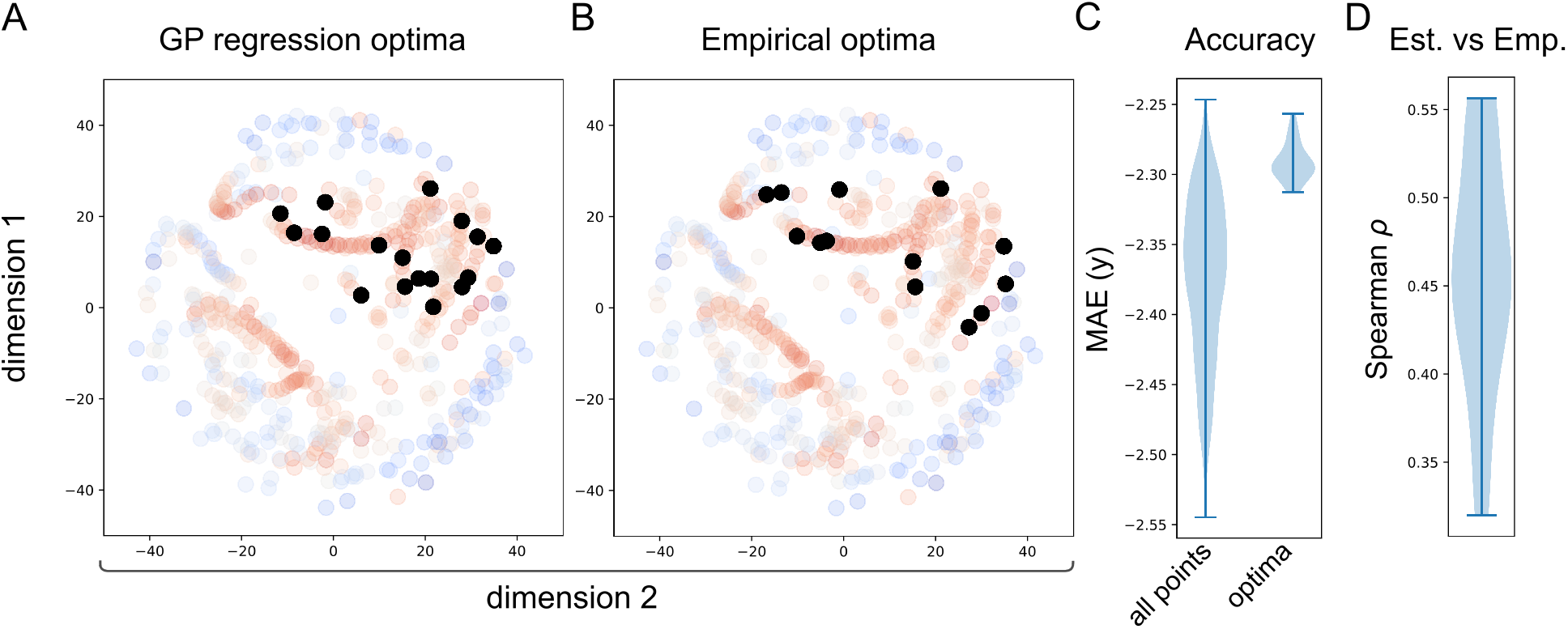
Performance of the optimization across different random starting conditions. For computational efficiency, only 20 iterations of active sampling were performed. Black dots represent optima of the 20 iterations based on A) the highest accuracy estimated using the GP model and B) the actual sampled points. C) Range of negative mean absolute error for the optima versus negative mean absolute errors across the whole space. D) correlations between actual and estimated accuracies across the whole space for the 20 replications.

## 4 Discussion

In this paper, we have established that active sampling can be used to map out a low-dimensional space of the multiverse of analytic approaches allowing the processing pipelines with higher accuracy to be identified in an efficient manner. We focused on a question with a known ground truth, predicting brain age from resting state functional connectivity data. Since efficient exploratory research is critical for neuroimaging if it is to become a mature scientific discipline, our multiverse approach is a critical tool that balances the need for rapid discovery with analytic rigor in a manner that is highly cost-efficient.

Our application of active sampling to predict age from functional connectivity is an illustrative example. We used a recently released dataset which was pre-processed using both motion regression and global signal regression [30]. Our main aim was to showcase active sampling on a space of analysis approaches, rather than identify (the) optimal combination(s) of fMRI head motion correction, functional connectome threshold and graph theoretical method for age prediction. Nevertheless, it remains interesting to consider the approaches selected as optimal by the GP regression. By repeating the active learning method 20 times, we see substantial consistency in processing steps across the selected optima. The motion regression approach consistently outperforms the global signal regression; lower, but not the lowest sparsities were also favored using a range of both simple and complex graph theoretical metrics, with betweenness centrality selected most frequently. These results are important because global signal regression is one of the most debated fMRI processing steps, with many arguments proposed both for and against its inclusion in processing pipelines [31, 32]. With respect to thresholding we observed that most of the optima had a higher threshold. This is in line with previous research, that observed that connections with lower edge weights (i.e., correlation) are more likely to be spurious suggesting that connectomes thresholded to lower densities might be less affected by noise [10, 30]. Finally, betweenness centrality had previously been found to perform well in network neuroimaging applications, including in machine learning applications [33].

Our multiverse approach is highly generalisable and so can easily be expanded to consider different analysis approaches and pre-processing techniques. Our method is not limited to graph theory and can be applied to any set of heterogeneous techniques in neuroimaging that can be evaluated by a common target measure, as is the case of different machine learning pipelines that try to minimize a cost function. For example, a similar approach could be used with full pre-processing pipelines used in (f)MRI and potentially integrated with automated pipelines such as fMRIPrep [34], to allow controlled, efficient exploration of a much wider range of analyses. We note, that given that the analysis space is based on variability across individuals, it does not require that the analysis approach results in data of the same format; it is possible to combine univariate and multivariate analyses (e.g., single regions, or every voxel or vertex measured), and even potentially different modalities, allowing for the multiverse to cover a very heterogeneous collection of approaches.

In the current paper, the analysis space was developed from a subset of the whole participant group; however, this need not be the case. A predefined space can be constructed using an existing dataset, and subsequently applied, with minimal computational cost, to different datasets. For example, large open datasets such as the Human Connectome Project or UK Biobank could be used to define analysis spaces which can then be applied to smaller, e.g., clinical datasets. This would mirror the strategy taken with many deep learning approaches, which are computationally expensive to train but not to apply to new data.

Performing multiverse analyses has the potential for increasing the generalizability of results, similarly to other approaches (e.g., [35]). As recently revisited by Yarkoni [36], when interpreting findings, we often go (both statistically and verbally) far beyond what is justified by the restricted nature of the data and analyses performed. Taking a multiverse approach explicitly tests the generalisability of the analyses: indeed, the GP regression model explicitly quantifies the relationships between analysis approaches, in the low-dimensional space. This can clarify how specific or general a given finding is across all of the approaches. The efficiency of the sampling of the space also ensures that the same data is only used a limited number of times, reducing the problems inherent in sequential analyses in terms of overfitting. In the extreme, in order to maximize generalisability, it is possible to perform each iteration of the active sampling on a different subset of participants who are not then reused; as such, each suggestion from the Bayesian optimization for the next point to be sampled would involve out-of-sample prediction.

Similarly to previous work using Bayesian optimization for the navigation of pre-defined experimental spaces [37, 38], the method presented here can help improve the poor reproducibility present across much of (neuro)science. Sequential analysis as applied here is highly formalized, quantifiable and controllable, and as such, it can be readily combined with pre-registration [39]. Similarly, the route and samples taken by the analysis make it possible to deduce what the hypothesis (encoded as the target function of the optimization algorithm) was at time of testing. If a different target function was selected, then the algorithm would have taken a different route through the analysis space (see [39]. This means that questionable research practices such as SHARKing may be more difficult to pursue [40].

As with any analysis approach, using active sampling methodologies comes with inherent trade-offs. Most notably, for more exploitative problems, where the optimal analysis approach is known (or approximately known) a priori or highly theoretically constrained, then the additional costs (in terms of sequential analysis affecting statistical power and computational burden) are a serious limitation. The optimization algorithm finding local minima resulting in poor overall performance is another potential limitation; this will depend heavily on the acquisition function including the type used and hyperparameters controlling exploration and exploitation as well as decisions regarding the GP regression and types of kernels used to model the low-dimensional space. A related issue is the creation of the low-dimensional space itself; this will inevitably involve a trade-off between capturing relevant variance and creating a relatively simple search space, with few dimensions. We show that the search space is coherent (in terms of the placement of similar analysis approaches near each other - Figure 2) and the GP regression is able to capture regularities in the space efficiently (Figure 3). However, for other problems, e.g., involving lower signal to noise, more heterogeneous variability across individuals, or more heterogeneous analysis approaches building a compact search space may be more challenging. Future work is needed to find the most useful acquisition function, GP regression and search spaces for applying active sampling approaches to multiverse analyses.

## Conclusion

We have presented a method for efficiently exploring outcomes across multiple analyses by building a low-dimensional space capturing the similarity of analysis approaches and subsequently exploring it using active learning. This enables maintaining the sensitivity of inferential statistics and generalizability to out of sample data while mapping out the multiverse of analysis approaches. Although we have illustrated the potential of this analysis by using FC data to predict brain age, this approach could be applied to different neuroimaging problems (both functional and structural) or indeed other types of data.

## 5 Availability of Code and Data

All code used for analyses and figure generation is available on GitHub (https://github.com/Mind-the-Pineapple/into-the-multiverse) and can be run using Colab. All data was previously released by Váša et al. [6] and is available on Figshare (https://doi.org/10.6084/m9.figshare.11551602).

## 6 Acknowledgments

R.L was funded by the MRC (Ref: MR/R005370/1) and by the Wellcome/EPSRC Centre for Medical Engineering (Ref: WT 203148/Z/16/Z); J.D. is funded by the King’s College London Imperial College London EPSRC Centre for Doctoral Training in Medical Imaging (EP/L015226/1).The authors would also like to acknowledge support from the Data to Early Diagnosis and Precision Medicine Industrial Strategy Challenge Fund, UK Research and Innovation (UKRI).

## 7 Supplementary Material

**Figure S1:**
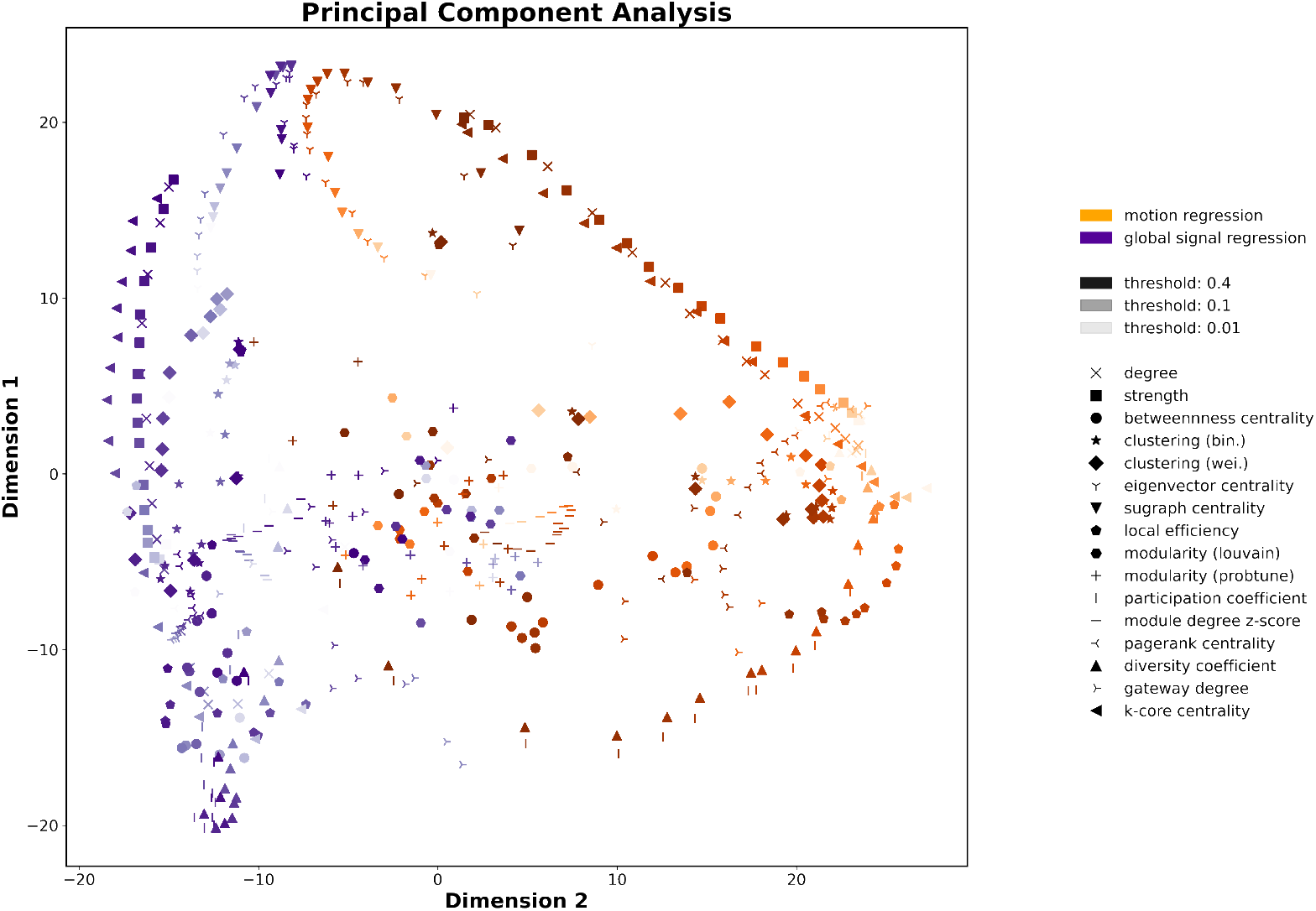
Dimensionality reduction using Principal Component Analysis (PCA). Similar to Figure 2, each point represents a different pipeline.

**Figure S2:**
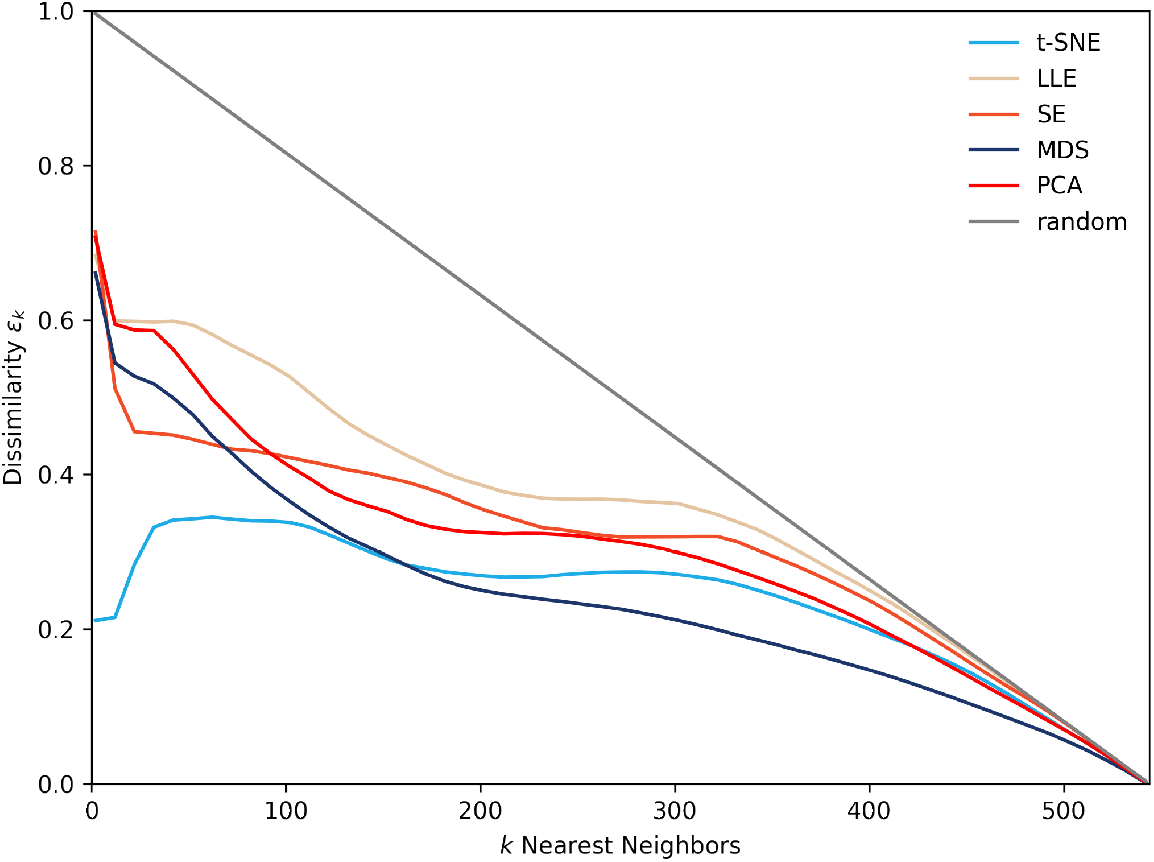
Dissimilarity score of the different low-dimensional embedding. The dissimilarity value quantifies how much the neighborhoods were maintained from the original to the reduced space; this was used to assess how much the different embedding algorithms maintained the topological information present in the original space. The dissimilarity score can assume values between 0 and 1; while small values mean that the neighborhood was similar after the embedding, values close to one signifies that the neighborhood was not conserved after the embedding. LLE, SE and PCA showed high dissimilarity values compared to MDS. Although t-SNE and UMAP showed a good similarity for a small number of neighbors, the MDS algorithm was able to keep the closest similarity for bigger neighborhoods. The gray line represents the dissimilarity score that would be achieved if all the neighbors would be randomly shuffled. This nicely illustrates that for very large number of clusters, the similarity scores are very close to the random performance.

## Notes

### Competing Interest Statement

The authors have declared no competing interest.

## References

[1] Joshua Carp. On the plurality of (methodological) worlds: Estimating the analytic flexibility of fmri experiments. Frontiers in neuroscience, 6:149, 10 2012.

[2] Danilo Bzdok and B T Thomas Yeo. Inference in the age of big data: Future perspectives on neuroscience. Neuroimage, 155:549–564, 07 2017.

[3] Rotem Botvinik-Nezer, Felix Holzmeister, Colin F Camerer, Anna Dreber, Juergen Huber, Magnus Johannesson, Michael Kirchler, Roni Iwanir, Jeanette A Mumford, R Alison Adcock, et al. Variability in the analysis of a single neuroimaging dataset by many teams. Nature, pages 1–7, 2020.

[4] Sara Steegen, Francis Tuerlinckx, Andrew Gelman, and Wolf Vanpaemel. Increasing Transparency Through a Multiverse Analysis. Perspect. Psychol. Sci., 11(5):702–712, sep 2016.

[5] Burr Settles. Active learning literature survey. Technical report, University of Wisconsin-Madison Department of Computer Sciences, 2009.

[6] František Váša, Rafael Romero-Garcia, Manfred G Kitzbichler, Jakob Seidlitz, Kirstie J Whitaker, Matilde M Vaghi, Prantik Kundu, Ameera X Patel, Peter Fonagy, Raymond J Dolan, Peter B Jones, Ian M Goodyer, the NSPN Consortium, Petra E Vértes, and Edward T Bullmore. Conservative and disruptive modes of adolescent change in human brain functional connectivity. Proc. Natl. Acad. Sci. U. S. A., 117 (6):3248–3253, 2020.

[7] Ed Bullmore and Olaf Sporns. Complex brain networks: graph theoretical analysis of structural and functional systems. Nat. Rev. Neurosci., 10(3):186–198, 2009.

[8] Martijn P. van den Heuvel and Hilleke E. Hulshoff Pol. Exploring the brain network: A review on resting-state fMRI functional connectivity. Eur. Neuropsychopharmacol., 20(8):519–534, aug 2010.

[9] Alex Fornito, Andrew Zalesky, and Michael Breakspear. Graph analysis of the human connectome: Promise, progress, and pitfalls. Neuroimage, 80:426–444, 2013.

[10] Martijn P. van den Heuvel, Siemon C. de Lange, Andrew Zalesky, Caio Seguin, B.T. Thomas Yeo, and Ruben Schmidt. Proportional thresholding in resting-state fMRI functional connectivity networks and consequences for patient-control connectome studies: Issues and recommendations. Neuroimage, 152:437–449, 2017.

[11] Mikail Rubinov. Constraints and spandrels of interareal connectomes. Nat. Commun., 7:1–11, 2016.

[12] David J Phillips, Alec Mcglaughlin, David Ruth, Leah R Jager, and Anja Soldan. NeuroImage : Clinical Graph theoretic analysis of structural connectivity across the spectrum of Alzheimer 3 s disease : The importance of graph creation methods. NeuroImage Clin., 7:377–390, 2015.

[13] James H Cole and Katja Franke. Predicting Age Using Neuroimaging: Innovative Brain Ageing Biomarkers. Trends Neurosci., 40(12):681–690, dec 2017.

[14] Tobias Kaufmann, Dennis van der Meer, Nhat Trung Doan, Emanuel Schwarz, Martina J Lund, Ingrid Agartz, Dag Alnæs, Deanna M Barch, Ramona Baur-Streubel, Alessandro Bertolino, and et al. Common brain disorders are associated with heritable patterns of apparent aging of the brain. Nat Neurosci, 22(10):1617–1623, 10 2019.

[15] J H Cole, S J Ritchie, M E Bastin, M C Valdés Hernández, S Muñoz Maniega, N Royle, J Corley, A Pattie, and S E Harris. Brain age predicts mortality. Nat. Publ. Gr., 23(5):1385–1392, 2017.

[16] Marc-Andre Schulz, B T Thomas Yeo, Joshua T Vogelstein, Janaina Mourao-Miranada, Jakob N Kather, Konrad Kording, Blake Richards, and Danilo Bzdok. Different scaling of linear models and deep learning in ukbiobank brain images versus machine-learning datasets. Nat Commun, 11(1):4238, 08 2020.

[17] Linda Geerligs, Remco J Renken, Emi Saliasi, Natasha M Maurits, and Monicque M Lorist. A brain-wide study of age-related changes in functional connectivity. Cerebral cortex, 25(7):1987–1999, 2015.

[18] Ricardo Pio Monti, Alex Gibberd, Sandipan Roy, Matthew Nunes, Romy Lorenz, Robert Leech, Takeshi Ogawa, Motoaki Kawanabe, and Aapo Hyvärinen. Interpretable brain age prediction using linear latent variable models of functional connectivity. Plos one, 15(6):e0232296, 2020.

[19] Matthew F. Glasser, Timothy S. Coalson, Emma C. Robinson, Carl D. Hacker, John Harwell, Essa Yacoub, Kamil Ugurbil, Jesper Andersson, Christian F. Beckmann, Mark Jenkinson, Stephen M. Smith, and David C. Van Essen. A multi-modal parcellation of human cerebral cortex. Nature, pages 1–11, 2016.

[20] Mikail Rubinov and Olaf Sporns. Complex network measures of brain connectivity: Uses and interpretations. Neuroimage, 52(3):1059–1069, 2010.

[21] B. T. Thomas Yeo, Fenna M. Krienen, Jorge Sepulcre, Mert R. Sabuncu, Danial Lashkari, Marisa Hollinshead, Joshua L. Roffman, Jordan W. Smoller, Lilla Zöllei, Jonathan R. Polimeni, Bruce Fischl, Hesheng Liu, and Randy L. Buckner. The organization of the human cerebral cortex estimated by intrinsic functional connectivity. J. Neurophysiol., 106(3), 2011.

[22] Sam T Roweis and Lawrence K Saul. Nonlinear dimensionality reduction by locally linear embedding. science, 290(5500):2323–2326, 2000.

[23] Mikhail Belkin and Partha Niyogi. Laplacian eigenmaps for dimensionality reduction and data representation. Neural computation, 15(6):1373–1396, 2003.

[24] Laurens van der Maaten and Geoffrey Hinton. Visualizing data using t-sne. Journal of machine learning research, 9(Nov):2579–2605, 2008.

[25] Joseph B Kruskal. Nonmetric multidimensional scaling: a numerical method. Psychometrika, 29(2):115–129, 1964.

[26] Leland McInnes, John Healy, Nathaniel Saul, and Lukas Grossberger. Umap: Uniform manifold approximation and projection. The Journal of Open Source Software, 3(29):861, 2018.

[27] Bobak Shahriari, Kevin Swersky, Ziyu Wang, Ryan P Adams, and Nando De Freitas. Taking the Human Out of the Loop : A Review of Bayesian Optimization. Proc. IEEE, 104(1):148–175, 2016.

[28] F. Pedregosa, G. Varoquaux, A. Gramfort, V. Michel, B. Thirion, O. Grisel, M. Blondel, P. Prettenhofer, R. Weiss, V. Dubourg, J. Vanderplas, A. Passos, D. Cournapeau, M. Brucher, M. Perrot, and E. Duchesnay. Scikit-learn: Machine learning in Python. Journal of Machine Learning Research, 12:2825–2830, 2011.

[29] Fernando Nogueira. Bayesian Optimization: Open source constrained global optimization tool for Python, 2014–.

[30] František Váša, Edward T. Bullmore, and Ameera X. Patel. Probabilistic thresholding of functional connectomes: Application to schizophrenia. Neuroimage, 172(May 2018):326–340, 2018.

[31] Kevin Murphy and Michael D. Fox. Towards a consensus regarding global signal regression for resting state functional connectivity MRI. Neuroimage, 154(November 2016):169–173, 2017.

[32] Jingwei Li, Ru Kong, Csaba Orban, Yanrui Tan, Nanbo Sun, Avram J Holmes, Mert R Sabuncu, Tian Ge, and B T Thomas Yeo. Global signal regression strengthens association between resting-state functional connectivity and behavior. Neuroimage, 196(April):126–141, 2019.

[33] Erik D Fagerholm, Peter J Hellyer, Gregory Scott, Robert Leech, and David J Sharp. Disconnection of network hubs and cognitive impairment after traumatic brain injury. Brain, pages 1696–1709, 2015.

[34] Oscar Esteban, Christopher J Markiewicz, Ross W Blair, Craig A Moodie, A Ilkay Isik, Asier Erramuzpe, James D Kent, Mathias Goncalves, Elizabeth Dupre, Madeleine Snyder, Hiroyuki Oya, Satrajit S Ghosh, Jessey Wright, Joke Durnez, Russell A Poldrack, and Krzysztof J Gorgolewski. fMRIPrep: a robust preprocessing pipeline for functional MRI. Nat. Methods, 16(January), 2019.

[35] Beth Baribault, Chris Donkin, Daniel R Little, Jennifer S Trueblood, Zita Oravecz, Don van Ravenzwaaij, Corey N White, Paul De Boeck, and Joachim Vandekerckhove. Metastudies for robust tests of theory. Proc Natl Acad Sci U S A, 115(11):2607–2612, 03 2018.

[36] Tal Yarkoni. The generalizability crisis. PsyArXiv, 2019.

[37] Romy Lorenz, Ricardo Pio Monti, Inês R. Violante, Christoforos Anagnostopoulos, Aldo A. Faisal, Giovanni Montana, and Robert Leech. The Automatic Neuroscientist: A framework for optimizing experimental design with closed-loop real-time fMRI. Neuroimage, 129:320–334, 2016.

[38] Romy Lorenz, Ines R Violante, Ricardo Pio Monti, Giovanni Montana, and Adam Hampshire. Dissociating frontoparietal brain networks with neuroadaptive Bayesian optimization. Nat. Commun., 2018.

[39] Romy Lorenz, Adam Hampshire, and Robert Leech. Neuroadaptive Bayesian Optimization and Hypothesis Testing. Trends Cogn. Sci., 21(3):155–167, 2017.

[40] Russell A Poldrack, Chris I Baker, Joke Durnez, Krzysztof J Gorgolewski, Paul M Matthews, Marcus R Munafò, Thomas E Nichols, Jean-baptiste Poline, Edward Vul, and Tal Yarkoni. Scanning the horizon : towards transparent and reproducible neuroimaging research. Nat. Publ. Gr., 2017.

